# DeepDiff-SHAP: Interpretable deep learning for subgroup-specific causal inference using conditional SHAP

**DOI:** 10.1101/2025.08.14.670190

**Authors:** Aditya Sriram, Soyeon Kim, Joseph A Carcillo, Hyun Jung Park

## Abstract

Precision medicine aims to tailor healthcare strategies to individual differences in genetic, clinical, and environmental factors. However, identifying subgroup-specific causal relationships in complex biomedical data remains a ma*j*or challenge, especially when standard causal inference methods average over population heterogeneity. We introduce DeepDiff-SHAP, a novel framework that combines regression-based and deep learning-based differential causal inference to detect changes in causal relationships across patient subgroups. DeepDiff-SHAP integrates SHapley Additive exPlanations (SHAP) to estimate conditional dependencies and perform nonlinear differential causal inference in a principled, interpretable manner. Applying DeepDiff-SHAP to two population-scale datasets, the CDC Diabetes Health Indicators Dataset and a UK Biobank sepsis cohort stratified by hypertension, we identified clinically meaningful and subgroup-specific causal changes in relationships between features such as age, general health, alkaline phosphatase, and cholesterol. Our results demonstrate that deep learning enhances sensitivity to complex interaction patterns overlooked by linear models, providing new biological insights into disease progression and comorbidity-specific risk mechanisms. DeepDiff-SHAP offers a scalable and interpretable solution to uncover individualized causal pathways, advancing the goal of truly personalized medicine.

## 1. Introduction

Precision medicine is transforming biomedical research and clinical care by shifting the focus from one-size-fits-all treatments to strategies tailored to individual differences in genetic, environmental, and lifestyle factors. As precision medicine continues to gain traction, there is a growing need for analytical methods that can identify subgroup-specific risk factors that either influence disease risk or therapeutic responses differently across distinct populations, or differently across multiple states within the same population. Examples include genetic variants such as *rs11673407* in the fucosyltransferase 3 gene (*FUT3*) elevating cardiovascular risk in men but not in women^1^, or nine potentially protective and 25 harmful metabolic biomarkers predicting future incidence of type 2 diabetes^2^. Traditional causal inference frameworks, however, typically estimate average treatment or exposure effects across a global population grouping^3-5^, which can obscure possible nuanced, group-specific mechanisms and lead to ineffective or even harmful interventions in underrepresented subgroups.

Recently, a method proposed by Belyaeva et al. called Differential Causal Inference (DCI), established a principled approach to detect differences in causal effects across groups by directly comparing the strength of variable-outcome relationships in one subgroup versus another^6^. DCI is methodologically distinct from simply performing causal inference separately in two groups and comparing the results. In a standard two-group approach, causal models are estimated independently for each group (e.g., diseased and healthy), and differences in effect estimates are compared post hoc. However, this approach does not account for estimation variance or the statistical significance of the differences, potentially leading to spurious findings or uncertainty in drawing conclusions. In contrast, DCI directly models and tests the difference in causal effects between groups as the primary quantity of interest within a unified framework that allows for more robust inference. This enables the identification of subgroup-specific causal mechanisms while controlling for variability and potential biases. Furthermore, the naive approach of fitting models separately in subgroups, especially in high-dimensional settings, can lead to unstable estimates due to small sample sizes or overfitting. DCI’s framework borrows strength across groups through *j*oint modeling or shared representations, improving estimation accuracy and interpretability. This enables researchers and clinicians to uncover risk factors that are uniquely relevant to particular population subgroups, such as non-responders to immunotherapy^7^ or patients with treatment-resistant depression^8^, thus advancing the development of more precise and targeted interventions.

Despite its advantage as being the only current differential causal inference method, DCI is based on regression-based framework and thus limited in its ability to capture the complex mechanisms underlying disease heterogeneity. Disease heterogeneity arises from complex, multilayered biological processes that involve nonlinear interactions among genetic, epigenetic, and environmental factors. Regression models, which rely on additive and linear assumptions, are insufficient to capture these complex dependencies, potentially overlooking key disease-driving mechanisms.

To model complex biological and clinical systems, which are governed by intricate, nonlinear interactions among molecular signals (e.g., gene expression, DNA methylation)^9,10^, environmental exposures (e.g., smoking, pollution)^11,12^, and clinical variables (e.g., comorbidities, treatment history)^13,14^, we will extend regression-based DCI using deep learning (DL). With its multilayered architecture and nonlinear activation functions, DL can effectively learn hierarchical feature representations that capture subtle, high-order dependencies between inputs. It has already demonstrated success in a variety of biomedical tasks including disease risk prediction^15, 16^ and image-based diagnostics^17,18^. This makes deep learning particularly well-suited for extending DCI for contexts in which causal effects are likely to vary not *j*ust in magnitude, but in functional form, across subgroups.

Specifically, between input feature A to B conditioning on *C* (**Fig. 1A**), we will extend the well-established regression-based three-step approach of DCI ^6^ (**Fig. 1B**) using DL architecture. While this approach evaluates model parameters for the conditional probability of each variable pair between conditions, performing these steps requires conditional probabilities, e.g., *P(A*|*B,S)*, where Cis a conditioning set. In the regression setting, such conditional distributions have closed-form expressions based on covariances and are thus tractable. However, in DL settings, these conditional distributions become complex and high-dimensional. To address this, we propose a novel deep learning framework, **DeepDiff-SHAP**, that incorporates advanced SHapley Additive exPlanations (SHAP)^19^ to quantify differential conditional dependence between variables (**Fig. 1C**). SHAP has been adapted to estimate conditional expectations in supervised learning^20^, but its use for computing conditional dependencies between input variables, as required for formal causal inference, remains limited and underexplored. By adapting SHAP to contrast feature contributions of certain variables, while conditioning on the shared, high-dimensional covariate space that exists in most biomedical data, our method enables robust and interpretable identification of variables whose causal effects differ across groups. This approach directly targets the core ob*j*ective of DCI and offers a scalable, principled solution for uncovering subgroup-specific causal mechanisms in complex biomedical data.

**Figure 1.**
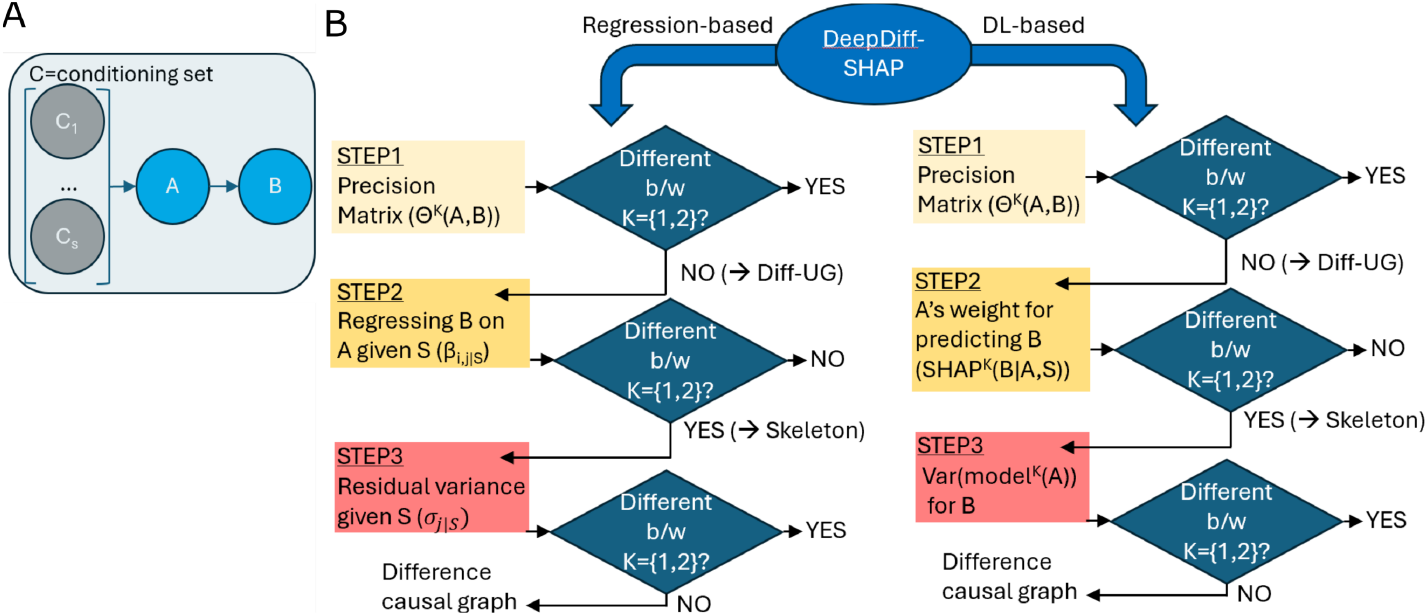
(A) Causal relationship from A to B to test for the difference based on the conditioning set C={C1,…,Cs}. (B) Algorithm of DeepDiff-SHAP with the regression and the deep learning components.

## 2. DeepDiff-SHAP: Overview and Algorithm Design

We introduce DeepDiff-SHAP as a principled, statistically sound three-algorithm approach rooted in regression and DL frameworks for causal structure changes between two states. DeepDiff-SHAP works without having to fully reconstruct each underlying network in a dataset. Step 1 involves identifying candidate nodes and edges where the dependency structure differs between states, based on changes in the precision matrix; in this case, the precision matrix is the pseudoinverse of the empirical covariance matrix. Since structural changes in a causal graph often result in shifts in the precision matrix, this step ensures a stable, high recall starting point for further evaluation and refinement. In Step 2, we test whether the strength of variable dependency relationships (i.e., how much one variable predicts another after conditioning on a third variable in the undirected graph) remains stable across the two states. If a relationship does not change, it is pruned from the candidate set, thus removing possible false positives identified by Step 1. Finally, in Step 3, we compute differences in residual (unexplained) variances calculated via deep neural networks (DNNs) to infer directionality between nodes; if there is statistical evidence of DNN residual invariance when conditioning on a set of nodes, it suggests that the node set contains the true causal parents related to the outcome.

### 2.1. Estimation of the difference undirected graph

To identify pairwise changes in conditional dependence structure between two states, we begin by estimating a **difference undirected graph (Δ-UG)** that aims to identify statistically significant evidence of interactions that vary between the two state-separated data groups. Let 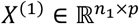 and 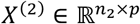 denote independent data groups from states 1 and 2, respectively. Each dataset is denoted by state-specific precision matrices 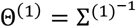 and 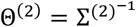, where *Σ*^*(k)*^is the covariance matrix of state *k*.

We implement a constraint-based approach that computes an edge-specific test statistic for each pair of variables (i,*j*) based on their estimated precision matrix entries. This step is adapted from the framework first formalized by Belyaeva et al^21^. Specifically, we estimate 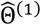 and 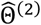 using the Moore–Penrose pseudoinverse of the empirical covariance matrices computed from *X*^(1)^ and *X*^(2)^ respectively. The test statistic for each pair (i,*j*) is defined as:

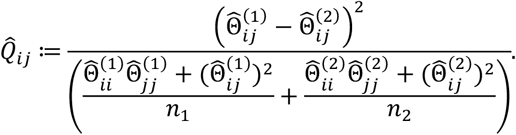

This statistic quantifies the squared difference in partial correlations between the two states, scaled by their estimated variances. Under the null hypothesis 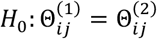, the statistic 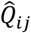 asymptotically follows a noncentral *F*-distribution^22,23^:

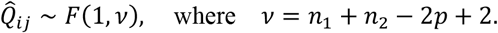

We compute p-values from this distribution using the cumulative noncentral *F* distribution function and define a significance threshold α ∈ [0,1]. The difference undirected graph Δis then constructed as:

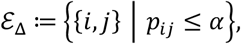

where *p*_i*j*_ is the *p*-value corresponding to 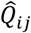. To reduce the downstream hypothesis testing burden in skeleton discovery and edge orientation, we define the set of conditioning nodes as:

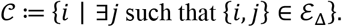

We limit the nodes included in the conditioning set to the nodes involved in the edges of the difference undirected graph, Δ-UG. We block the inclusion of any additional nodes i.e. marginal or conditional distributions differing across states leading to node inclusion in the conditioning set, resulting in a very strict edge inclusion step.

### 2.2. Skeleton discovery via SHAP-based conditional invariance testing

In the second step of our method, we further prune the initial undirected difference graph by testing for conditional invariance of cross-state feature dependencies. Specifically, for each edge (*i, j*) in the initial undirected difference graph, we assess whether the importance of feature *i* for predicting feature *j* remains invariant across states, and vice versa, after conditioning on subsets of features part of the conditioning set *S*.

We model each feature *i* as a function of its potential parent *j* and a conditioning set *S* ⊆ 𝒞{*i, j*}, where 𝒞 denotes the set of conditioning nodes obtained from Step 1. For each direction, we train two multilayer perceptron regressors 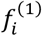 and 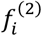 on the two datasets *X*^(1)^ and *X*^(2)^ to predict *X*_i_ using *X*_*j*_ and *S*.

To assess the contribution of *X*_*j*_ to the prediction of *X*_*i*_ in each state, we compute conditional SHAP values using KernelSHAP with a fixed background distribution that isolates the effect of *X*_*j*_ given *S*^24^. Specifically, we calculate:

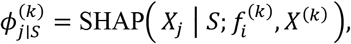

where *k* ∈ {1,2} and 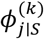 denotes the absolute SHAP values across test samples. These values are then compared between states using a normalized squared difference statistic:

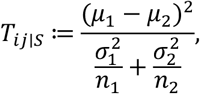

where *μ*_*k*_ and 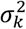 are the mean and variance of the SHAP values in state *k*, and *n*_*k*_ is the number of SHAP samples. A two-sided *p*-value is derived from the noncentral *F*-distribution with degrees of freedom ν = *n*_1_ + *n*_2_ − 2 − 2|*S*|.

For each ordered pair (*i* ←*j* and *j* ← *i*) we test for SHAP heterogeneity across states conditional on the set of conditioning nodes. When *p*> α, we fail to re*j*ect the null hypothesis of conditional invariance across states and remove the corresponding edge from the undirected difference graph. When *p*< *α*, we reject the null hypothesis of conditional invariance, and we include the edge. This process is repeated across all conditioning sets up to a specified maximum size *r*_max_. The remaining edges after this pruning step make up the edge difference skeleton used in the subsequent direction-orientation step for the leftover edges.

This SHAP-based invariance testing allows for nonlinear, model-flexible detection of asymmetric changes in feature relationships, extending the regression-based conditional independence tests to deep neural network models with a more structurally sound framework.

### 2.3 Edge orientation via invariance testing

In step 3, we orient edges obtained in step 2 (the pruned edge difference skeleton) to end up with a directed graph that captures differences in functional dependencies across the two states. We assume that for any node *j*, if the conditional variance of *X*_*j*_ given a set of likely parents Sis invariant across states, then Srepresents a valid set of parents for *j*. This principle is rooted in the theory that the functional form of the conditional distribution *P*(*X*_*j*_ | *X*_*S*_) should remain stable under invariance^25^.

For each node *j* in the graph, we test candidate parent sets S⊆ 𝒞 {*j*} of size *k* = 1, where 𝒞 is the set of conditioning nodes identified from the pruned edge difference skeleton from algorithm 2. We train DNNs to regress *X*_*j*_ on *X*_*S*_ separately in each state using a two-layer multilayer perceptron (MLP). The residual variance is estimated as:

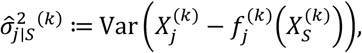

Where 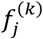 denotes the fitted DNN regressor in state *k* ∈ {1,2}, and 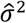 is computed on the training data. To assess whether the conditional variance differs significantly between states, we compute a two-sided test statistic based on the ratio of residual variances:

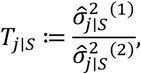

with a *p*-value computed using the noncentral *F*-distribution with degrees of freedom (*n*_1_ − |*S*|, *n*_2_ − |*S*|):

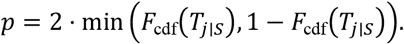

If the p-value exceeds a specified threshold α, we fail to re*j*ect the null hypothesis and conclude that the conditional variance is invariant, therefore accepting S as the parent set for *j*. The directed edges *i*→*j*for all *i* ∈ Sare added to the graph. We perform additional cycle and contradiction checks using transitive closure on the directed graph to prevent invalid orientations.

For any edges that remain unoriented after this test, we apply graph traversal rules to resolve directionality wherever a consistent path structure allows. For example, for any set of nodes in which *i*→ *node*_1_→ *node*_*x*_→ *j*, we orient *i*→*j*. This rationale is formalized by Meek and reviewed by Colombo et al^26,27^. The final output is a directed ad*j*acency matrix, a log of all the orientation decisions, and the set of edges that could not be oriented by algorithm 3’s invariance testing.

This DNN-based residual variance orientation strategy expands on the original DCI steps defined for regression-based models and leverages DL via a DNN framework to capture potentially nonlinear predictive structure, and tests whether this structure is preserved across states; this allows for stronger performance of causal inference in a model-independent way^21^. Our model eliminates the assumption of linear-Gaussian data by utilizing DNN-based predictions for variable dependencies.

For results mentioned in this paper, DeepDiff-SHAP was initialized with the following parameters: α_ug_ = 0.005, α_*skeleton*_ = 0.3, α_*orient*_ = 0.001 corresponding to the threshold levels for each of the three steps of the algorithm. Conditioning set size was set to 1 for (maximum set size is 2, range from 0 to 2; a higher conditioning set size leads to sparser causal network graphs).

## 3 Results

### 3.1 DeepDiff-SHAP reveals nonlinear subgroup-specific causal structures in diabetes populations

To investigate changes in causal relationships associated with chronic disease, we ran the regression-based DCI and DL-based DeepDiff-SHAP to the 2014 Centers for Disease Control and Prevention (CDC) Diabetes Health Indicators Dataset (9). This dataset includes health status, behavior, and access-to-care survey data for 253,680 individuals, among whom 39,977 were diagnosed as diabetic or prediabetic and 213,703 were not diagnosed with diabetes. Using the regression-based DCI module of DeepDiff-SHAP, which follows the original DCI framework, we detected 68 feature pairs with differential associations between the diabetic and non-diabetic groups. Many of these association changes involved well-known demographic confounders such as age and sex, which influence a wide range of lifestyle and health indicators. For instance, age was differentially associated with features like General Health (GenHlth), Stroke, Smoker (yes/no), and Access to Health Care (AnyHealthcare), while sex was associated with differences in Income, Smoking status, and BMI (**Fig. 2A**). 12 of the 68 differential associations (17.64%) were attributed to underlying causal relationship changes by the regression-based model. Among these, only 2 (16.66%) involved age or sex as causal variables, despite their central role in shaping health outcomes. For example, while it is clinically plausible that age impacts general health status differently in people with and without diabetes, this causal difference was not detected by the regression model. In contrast, the deep learning module of DeepDiff-SHAP (**Fig. 2B**) identified this expected causal difference from Age to GenHlth, as well as a differential causal effect of Age on High Cholesterol (HighChol). Similarly, while the regression-based model did not attribute any of the sex-related association changes, such as those from Sex to Heart Disease or Heart Attack (HeartDiseaseAttack), Smoking status, and BMI, to changes in causal relationships, the DL-based approach successfully identified all these as differential causal effects between the diabetes and non-diabetes groups.

**Figure 2.**
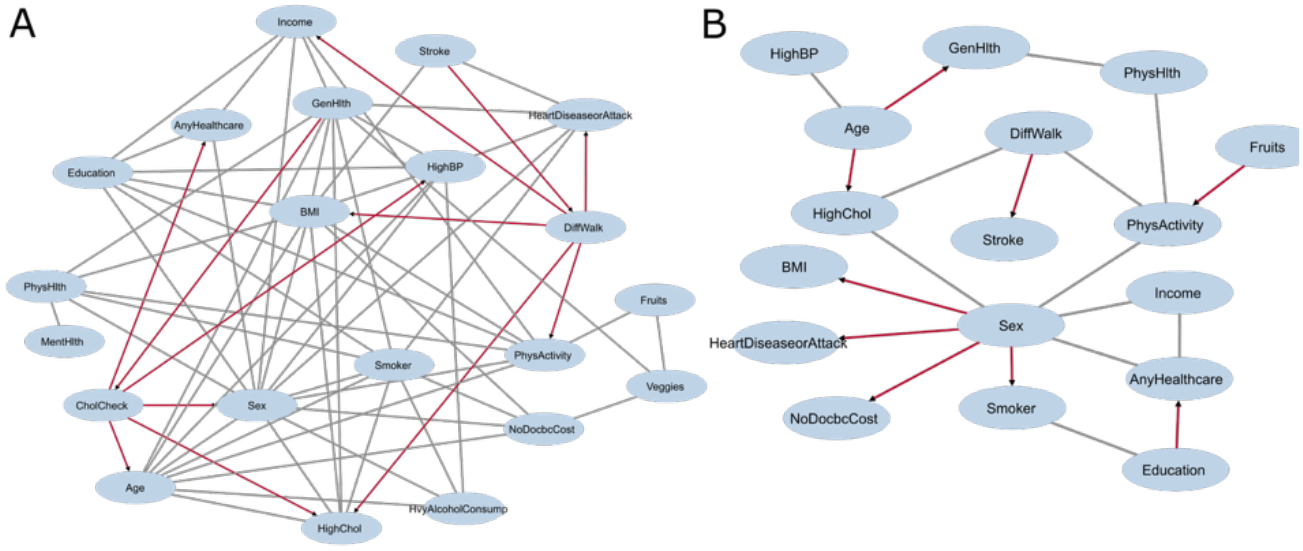
DCI graph on the diabetes data as identified by (A) the regression-based and (B) deep-learning-based DeepDiff-SHAP. Gray lines represent the association changes that are not attributed as differential causal relationship and red lines represent the differential causal relationships.

We note the importance of these particular causal relationships in the DeepDiff-SHAP network graph, Age to High Cholesterol and Sex to Coronary Heart Disease or Myocardial Infarction, both which have been previously mentioned in the context of type 2 diabetes being associated with women having a greater risk of cardiovascular disease, as well as studies implicating earlier cardiovascular disease events in women with T2D compared to men^28-30^. These findings suggest that deep learning-based causal inference can uncover subtle and nonlinear changes in causal structure that may be missed by linear models. Notably, the regression-based DCI model identified some relationships that appear potentially misleading. As an example, regression-based DCI attributed a causal difference in the relationship from Stroke to Difficulty Walking, which is more plausibly explained in the reverse direction or mediated by baseline functional impairments in diabetic individuals. Altogether, these results both highlight the limitations of traditional regression-based causal inference in detecting meaningful shifts in causal mechanisms between disease subpopulations as well as showcase how our deep learning framework enables more nuanced detection of differential causal structures, supporting its utility in understanding the biological and behavioral heterogeneity of chronic diseases such as diabetes.

### 3.2 Uncovering comorbidity-specific mechanisms in sepsis through causal inference

To investigate how chronic comorbidities modulate causal relationships in sepsis, we applied DeepDiff-SHAP to a subset of the UK Biobank comprising 3,181 individuals diagnosed with sepsis, stratified by hypertension status (hypertension: n = 2,669; no hypertension: n = 512). Sepsis remains a leading cause of morbidity and mortality in adults and children, yet its clinical progression is strongly influenced by pre-existing conditions such as hypertension, a factor often overlooked in risk modeling^31^. We identified sepsis cases using ICD-10 codes (e.g., A40.x, A41.x, B37.7, O85), capturing a range of septicemia and related conditions. From the UK Biobank, we selected 42 variables spanning domains of cardiac-metabolic health, renal and liver function, inflammation, hormones, blood pressure, and pulmonary status. Using DeepDiff-SHAP’s regression-based module, we identified 18 associations that differed between hypertensive and non-hypertensive sepsis patients, of which 8 (44.44%) were attributed to shifts in causal relationships (**Fig. 3A**). Notably, these included altered causal links such as from urate to SHBG and Apolipoprotein B, total bilirubin to triglycerides, calcium to total protein, Apolipoprotein A1 to systolic blood pressure, and IGF-1 to age at hypertension diagnosis, suggesting distinct physiological pathways influenced by vascular stress. Importantly, the deep learning component of DeepDiff-SHAP uncovered an additional causal shift from alkaline phosphatase to total cholesterol that was not detectable via regression model (**Fig. 3B**), highlighting the added value of incorporating nonlinear modeling. Together, these findings demonstrate that DeepDiff-SHAP enables a comprehensive discovery of subgroup-specific causal mechanisms in sepsis by integrating both regression and deep learning frameworks.

**Figure 3.**
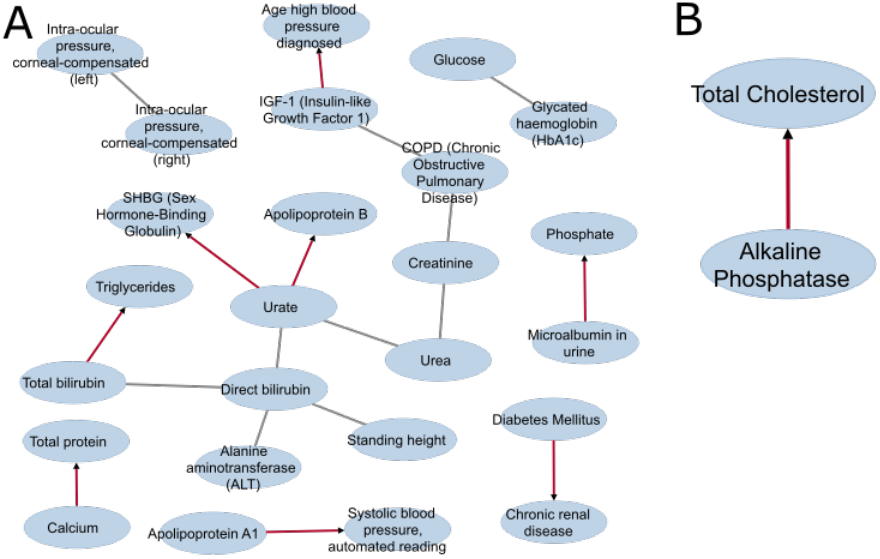
DCI graph on the sepsis data as identified by (A) the regression-based and (B) deep-learning-based DeepDiff-SHAP. Gray lines represent the association changes that are not attributed as differential causal relationship and red lines represent the differential causal relationships.

### 3.3 Benchmarking DeepDiff-SHAP against regression-based DCI: enhanced sensitivity to nonlinear causal differences

To benchmark performance, we compared DeepDiff-SHAP against DCI, the only existing differential causal inference method currently available, that is based entirely on regression modeling. Importantly, the regression-based component of DeepDiff-SHAP is mathematically equivalent to this original DCI method, as both frameworks follow the same three-step procedure: identifying structural differences, testing conditional invariance, and estimating directionality based on residual variance. Therefore, any observed differences in results between DeepDiff-SHAP’s deep learning (DL) module and the regression-based DCI can be attributed to the modeling framework itself, rather than the procedural design. To assess relative sensitivity to causal structure changes, we compared the number of association differences that were ultimately attributed to changes in causal strength (i.e., differential causal degrees). Since the comparison holds the inference pipeline constant while varying only the functional form (linear vs. nonlinear approach), this metric offers a controlled and clear evaluation of how effectively each approach detects meaningful causal differences. Between the diabetes (pre-diabetic and diabetic diagnosis) and healthy population groups in the CDC Diabetes dataset, the original DCI method identified 68 association changes, but only 12 (17.6%) were attributed to differences in causal degree. In contrast, DeepDiff-SHAP detected 19 association changes, with 9 (47.3%) resolved as true causal degree differences, demonstrating a significantly higher sensitivity in uncovering meaningful causal changes (P-value = 0.02). A similar pattern emerged in the sepsis dataset: DeepDiff-SHAP resolved 1 causal degree difference out of 1 association change, whereas the original DCI resolved 8 out of 18 association changes (P-value = 0.2). Although the weaker significance in the sepsis dataset may reflect the presence of only a single nonlinear association difference, these findings nonetheless underscore the improved sensitivity of DeepDiff-SHAP, particularly for detecting complex, nonlinear causal relationships.

## 4 Discussion

We introduce DeepDiff-SHAP as a method that builds on regression-based DCI with a deep learning-based differential causal inference framework. Our method’s modular design, split into three steps with flexible parameter initialization settings, allows researchers to systematically evaluate subgroup-specific differences in feature importance and directional relationships, creating a highly customizable approach to evaluating causal data structure. Additionally, the results observed from our comparative analysis across two unique patient datasets, the CDC Diabetes Health Indicators Dataset and a UK Biobank cohort of sepsis patients stratified by hypertension, demonstrate DeepDiff-SHAP’s ability to identify biologically meaningful and study-supported causal relationships that regression-only DCI may overlook.

DeepDiff-SHAP’s analysis of the Diabetes dataset offers novel insights into the differential roles of classical foundational variables such as sex and age. Diabetes is a chronic metabolic disease that compounds over time; individuals with diabetes are more likely to experience a steeper deterioration in general health as they age, compared to individuals without diabetes.

Additionally, aging itself brings about general decline in metabolic efficiency, immune function, and tissue repair, all of which can be exacerbated by the presence of diabetes. By leveraging DL’s multilayer modeling capacities, we reveal a synergistic effect between age and diabetes, where the impact of aging on general health is causally differential. In contrast, individuals without diabetes may experience a more gradual, less pronounced decline in general health with aging. Since such group-specific causal relationships like the effect of age on general health require capturing complex, higher-order interactions, our DL-based approach proved essential in uncovering them, underscoring its potential for advancing precision medicine in diabetes care.

Similarly, regarding the unique causal difference result between sepsis patients with and without hypertension, several studies have made the link between serum alkaline phosphatase (ALP) and coronary artery disease, with hypertension recognized as a well-known contributor to the development of coronary artery disease^32, 33^. Additional studies in humans and mice have linked ALP with elevated cholesterol levels, particularly in those with dyslipidemia, and observe that high ALP in dyslipidemia patients leads to hypertension and coronary heart disease^34,35^.

Despite the promise of DeepDiff-SHAP in identifying subgroup-specific differences in causal structure, our current framework has some limitations. First, the computational efficiency of the SHAP-based conditional invariance testing step remains a ma*j*or challenge. Specifically, estimating conditional SHAP values for each candidate variable pair across multiple conditioning subsets is computationally expensive, particularly for large-scale datasets with many variables and complex feature interactions. Second, our current approach uses an ad hoc restriction on the conditioning set to reduce complexity: we limit candidate conditioning variables to those involved in edges identified in the difference undirected graph (Δ-UG). While this helps avoid an exponential increase in the number of SHAP computations, it may miss subtler or higher-order conditional dependencies. This limitation can inherently bias results towards more prominent signal changes while underestimating other nuanced shifts in conditional structure. To address this, future iterations of DeepDiff-SHAP could benefit from dimensionality reduction techniques such as variable screening, supervised embedding, or attention-based feature selection before applying conditional SHAP. Finally, we will also investigate adaptive strategies for selecting conditioning sets that leverage measures such as mutual information or latent feature representations. The goal is to balance out computational feasibility and the capacity to capture potentially hidden dependencies. Methods such as autoencoder-based dimensionality reduction, graph neural network–derived embeddings, or Bayesian network–informed priors may help us develop principled ways to limit the conditioning space while still retaining the most relevant differential dependency structures.

As medicine increasingly moves toward personalized interventions, understanding how risk factors or biological pathways operate differently across patient subpopulations, such as those with or without comorbidities like diabetes or hypertension, is essential to avoid generalized solutions that can be ineffective or even harmful. Traditional regression-based methods average over heterogeneity, preventing subtle but important biological differences from being uncovered. Our approach, which integrates the theoretical rigor of differential causal inference with the interpretable power of deep learning and SHAP, allows researchers to discover changes in causal structure that vary with disease state, comorbidity, or population subgroup. These insights can directly inform the design of more precise diagnostic criteria, risk prediction tools, and treatment strategies, ultimately improving clinical outcomes by ensuring the appropriate interventions are delivered to the appropriate patients.

## 5. Acknowledgments

The CDC Diabetes Health Indicators dataset comes from the University of California, Irvine Machine Learning Repository and the CDC 2014 Annual Data report. UK Biobank data comes from the UK Biobank Resource under Application Number 83829. This research was supported in part by the University of Pittsburgh Center for Research Computing, RRID:SCR_022735, through the resources provided. Specifically, this work used the HTC cluster, which is supported by NIH award number S10OD028483.

## 6 Code Availability

DeepDiff-SHAP code and examples can be accessed at: https://github.com/ads303/DeepDiff-SHAP.

## References

1. Silander K, Alanne M, Kristiansson K, Saarela O, Ripatti S, Auro K, Karvanen J, Kulathinal S, Niemelä M, Ellonen P, Vartiainen E, Jousilahti P, Saarela J, Kuulasmaa K, Evans A, Perola M, Salomaa V, Peltonen L. Gender Differences in Genetic Risk Profiles for Cardiovascular Disease. PLOS ONE. 2008;3(10):e3615. doi: 10.1371/journal.pone.0003615.

2. Peddinti G, Cobb J, Yengo L, Froguel P, Kravić J, Balkau B, Tuomi T, Aittokallio T, Groop L. Early metabolic markers identify potential targets for the prevention of type 2 diabetes. Diabetologia. 2017;60(9):1740–50. doi: 10.1007/s00125-017-4325-0.

3. Sedgewick AJ, Buschur K, Shi I, Ramsey JD, Raghu VK, Manatakis DV, Zhang Y, Bon J, Chandra D, Karoleski C, Sciurba FC, Spirtes P, Glymour C, Benos PV. Mixed graphical models for integrative causal analysis with application to chronic lung disease diagnosis and prognosis. Bioinformatics 2019. p. 1204–12.

4. Zheng X, Aragam B, Ravikumar PK, Xing EP. DAGs with NO TEARS: Continuous Optimization for Structure Learning: Curran Associates, Inc.; 2018.

5. Moccia C, Moirano G, Popovic M, Pizzi C, Fariselli P, Richiardi L, Ekstrøm CT, Maule M. Machine learning in causal inference for epidemiology. European Journal of Epidemiology. 2024;39(10):1097–108. doi: 10.1007/s10654-024-01173-x.

6. Belyaeva A, Squires C, Uhler C. DCI: learning causal differences between gene regulatory networks. Bioinformatics 2021. p. 3067–9.

7. Davar D, Morrison RM, Dzutsev AK, Karunamurthy A, Chauvin J-M, Amatore F, Deutsch JS, Das Neves RX, Rodrigues RR, McCulloch JA, Wang H, Hartman DJ, Badger JH, Fernandes MR, Bai Y, Sun J, Cole AM, Aggarwal P, Fang JR, Deitrick C, Bao R, Duvvuri U, Sridharan SS, Kim SW, A. Choudry H, Holtzman MP, Pingpank JF, O’Toole JP, DeBlasio R, Jin Y, Ding Q, Gao W, Groetsch C, Pagliano O, Rose A, Urban C, Singh J, Divarkar P, Mauro D, Bobilev D, Wooldridge J, Krieg AM, Fury MG, Whiteaker JR, Zhao L, Paulovich AG, Najjar YG, Luke JJ, Kirkwood JM, Taube JM, Park HJ, Trinchieri G, Zarour HM. Neoadjuvant vidutolimod and nivolumab in high-risk resectable melanoma: A prospective phase II trial. Cancer Cell. 2024;42(11):1898-918.e12. doi: 10.1016/j.ccell.2024.10.007.

8. Ruberto VL, Jha MK, Murrough JW. Pharmacological Treatments for Patients with Treatment-Resistant Depression. Pharmaceuticals. 2020;13(6):116. PubMed PMID: doi:10.3390/ph13060116.

9. Olecka M, van Bömmel A, Best L, Haase M, Foerste S, Riege K, Dost T, Flor S, Witte OW, Franzenburg S. Nonlinear DNA methylation trajectories in aging male mice. Nature Communications. 2024;15(1):3074.

10. Zablotskii V, Gorobets O, Gorobets S, Polyakova T. Effects of Static and Low‐Frequency Magnetic Fields on Gene Expression. Journal of Magnetic Resonance Imaging. 2025.

11. Cheng W-C, Wong P-Y, Wu C-D, Cheng P-N, Lee P-C, Li C-Y. Non-linear association between long-term air pollution exposure and risk of metabolic dysfunction-associated steatotic liver disease. Environmental health and preventive medicine. 2024;29:7-.

12. Demateis D, Keller KP, Rojas‐Rueda D, Kioumourtzoglou MA, Wilson A. Penalized distributed lag interaction model: Air pollution, birth weight, and neighborhood vulnerability. Environmetrics. 2024;35(4):e2843.

13. Chesnaye NC, van Diepen M, Dekker F, Zoccali C, Jager KJ, Stel VS. Non-linear relationships in clinical research. Nephrology Dialysis Transplantation. 2025;40(2):244–54.

14. Zhang Y, Chen X, Wang Y, Tang Y, Zhang K, Wu J. Analysis of the nonlinear relationships between insulin resistance indicators such as LAP and TyG and depression, and population characteristics: a cross-sectional study. European Journal of Medical Research. 2025;30(1):513.

15. Oh TR. Integrating predictive modeling and causal inference for advancing medical science. Childhood Kidney Diseases. 2024;28(3):93–8.

16. Nkoy FL, Stone BL, Zhang Y, Luo G. A roadmap for using causal inference and machine learning to personalize asthma medication selection. JMIR Medical Informatics. 2024;12(1):e56572.

17. Deshpande S, Li Z, Kuleshov V. Multi-Modal Causal Inference with Deep Structural Equation Models. arXiv preprint 220309672. 2022.

18. Zang C, Wang H, Pei M, Liang W, editors. Discovering the real association: Multimodal causal reasoning in video question answering. Proceedings of the IEEE/CVF Conference on Computer Vision and Pattern Recognition; 2023.

19. Lundberg SM, Lee S-I. A Unified Approach to Interpreting Model Predictions. In: Guyon I, Luxburg UV, Bengio S, Wallach H, Fergus R, Vishwanathan S, Garnett R, editors. 2017.

20. Sundararajan M, Najmi A, editors. The many Shapley values for model explanation. International conference on machine learning; 2020: PMLR.

21. Belyaeva A, Squires C, Uhler C. DCI: learning causal differences between gene regulatory networks. Bioinformatics. 2021;37(18):3067–9. doi: 10.1093/bioinformatics/btab167. PubMed PMID: 33704425; PMCID: PMC9991896.

22. Lütkepohl H. New introduction to multiple time series analysis. Berlin;: New York: Springer; 2005. xxi, 764 p. p.

23. Yuhao Wang CS, Anastasiya Belyaeva, Caroline Uhler. Direct estimation of differences in causal graphs in Advances in Neural Information Processing Systems. NeurIPS 2018; Montreal: Advances in Neural information Processing Systems; 2018.

24. Scott M Lundberg S-IL. A Unified Approach to interpreting Model Predictions. International Conference on Neural Information Processing Systems plLong Beach, CA: Advances in Neural Information Processing Systems; 2017

25. Peters J, Bühlmann P, Meinshausen N. Causal Inference by using Invariant Prediction: Identification and Confidence Intervals. Journal of the Royal Statistical Society Series B: Statistical Methodology. 2016;78(5):947–1012. doi: 10.1111/rssb.12167.

26. Meek C. Causal inference and causal explanation with background knowledge. Proceedings of the Eleventh conference on Uncertainty in artificial intelligence; Montréal, Qué, Canada: Morgan Kaufmann Publishers Inc.; 1995. p. 403–10.

27. Diego Colombo MHM. Order-Independent Constraint-Based Causal Structure Learning. Journal of Machine Learning Research. 2014;15(116):3921—62.

28. Madonna R, Balistreri CR, De Rosa S, Muscoli S, Selvaggio S, Selvaggio G, Ferdinandy P, De Caterina R. Impact of Sex Differences and Diabetes on Coronary Atherosclerosis and Ischemic Heart Disease. J Clin Med. 2019;8(1). Epub 20190116. doi: 10.3390/jcm8010098. PubMed PMID: 30654523; PMCID: PMC6351940.

29. Yoshida Y, Chen Z, Fonseca VA, Mauvais-Jarvis F. Sex Differences in Cardiovascular Risk Associated With Prediabetes and Undiagnosed Diabetes. Am J Prev Med. 2023;65(5):854–62. Epub 20230514. doi: 10.1016/j.amepre.2023.05.011. PubMed PMID: 37192710.

30. Ballotari P, Venturelli F, Greci M, Giorgi Rossi P, Manicardi V. Sex Differences in the Effect of Type 2 Diabetes on Major Cardiovascular Diseases: Results from a Population-Based Study in Italy. Int J Endocrinol. 2017;2017:6039356. Epub 20170220. doi: 10.1155/2017/6039356. PubMed PMID: 28316624; PMCID: PMC5338069.

31. Ahlberg CD, Wallam S, Tirba LA, Itumba SN, Gorman L, Galiatsatos P. Linking Sepsis with chronic arterial hypertension, diabetes mellitus, and socioeconomic factors in the United States: A scoping review. Journal of Critical Care. 2023;77:154324. doi: 10.1016/j.jcrc.2023.154324.

32. Chen Y, Zhou ZF, Han JM, Jin X, Dong ZF, Liu L, Wang D, Ye TB, Yang BS, Zhang YP, Shen CX. Patients with comorbid coronary artery disease and hypertension: a cross-sectional study with data from the NHANES. Ann Transl Med. 2022;10(13):745. doi: 10.21037/atm-22-2766. PubMed PMID: 35957737; PMCID: PMC9358511.

33. Weber T, Lang I, Zweiker R, Horn S, Wenzel RR, Watschinger B, Slany J, Eber B, Roithinger FX, Metzler B. Hypertension and coronary artery disease: epidemiology, physiology, effects of treatment, and recommendations: A joint scientific statement from the Austrian Society of Cardiology and the Austrian Society of Hypertension. Wien Klin Wochenschr. 2016;128(13-14):467–79. Epub 20160609. doi: 10.1007/s00508-016-0998-5. PubMed PMID: 27278135.

34. Adamidis PS, Florentin M, Liberopoulos E, Koutsogianni AD, Anastasiou G, Liamis G, Milionis H, Barkas F. Association of Alkaline Phosphatase with Cardiovascular Disease in Patients with Dyslipidemia: A 6-Year Retrospective Study. J Cardiovasc Dev Dis. 2024;11(2). Epub 20240215. doi: 10.3390/jcdd11020060. PubMed PMID: 38392274; PMCID: PMC10889667.

35. Bessueille L, Kawtharany L, Quillard T, Goettsch C, Briolay A, Taraconat N, Balayssac S, Gilard V, Mebarek S, Peyruchaud O, Duboeuf F, Bouillot C, Pinkerton A, Mechtouff L, Buchet R, Hamade E, Zibara K, Fonta C, Canet-Soulas E, Millan JL, Magne D. Inhibition of alkaline phosphatase impairs dyslipidemia and protects mice from atherosclerosis. Transl Res. 2023;251:2–13. Epub 20220617. doi: 10.1016/j.trsl.2022.06.010. PubMed PMID: 35724933.

